# Prediction of quantitative function of artificially-designed protein from structural information

**DOI:** 10.1101/2025.04.10.648284

**Authors:** Ryosaku Ota, Masayuki Sakamoto, Wataru Aoki, Honda Naoki

## Abstract

Artificially designed proteins are widely used in applications such as optogenetics and biosensing. While experimental optimization of these proteins is effective, it is also costly and labor-intensive. To address this challenge, computational approaches have been developed, primarily relying on sequence-based features. However, protein function is inherently tied to its three-dimensional (3D) structure, and incorporating structural information could enable more accurate predictions and provide deeper biological interpretability. Here, we proposed a structure-based analysis framework called ‘Foldinsight’ for predicting protein functionalities. In our framework, we first predict protein structures from sequences using AlphaFold2 and then utilize these structures to predict protein properties. Since proteins vary in the number of atoms and lack direct atomic correspondence, we applied molecular field mapping, which captures the energy states surrounding a protein and converts them into fixed-length numerical vectors. This transformation enables the application of machine learning, allowing protein properties to be predicted from structure-derived features. Applying this framework to channelrhodopsin mutants, we achieved predictive performance comparable to sequence-based models. Additionally, our structure-based analysis successfully identified key structural regions contributing to functional differences, highlighting the advantage of incorporating structural data into predictive modeling.

## Introduction

Artificially designed proteins are engineered to enhance specific functionalities by introducing mutations into their natural proteins^1–3^. This approach has been widely applied in fields such as optogenetics and biosensing^4–7^. The process of optimizing these proteins involves a labor-intensive and iterative trial-and-error approach^8–11^. In this process, mutations are randomly introduced into the original protein sequence to improve desired characteristics^12–14^. Identifying mutations that significantly enhance functionality remains a major challenge due to the limited understanding of the relationship between a protein’s structure and its function.

To reduce the cost of screening, informatics approaches are increasingly favored^15–17^. By leveraging computational predictions, these approaches can complement wet experiments, providing a more efficient pathway. However, when creating prediction models, it is crucial to consider the physiological aspects of proteins. In principle, a protein’s function and performance are intrinsically linked to its three-dimensional (3D) structure^18–20^. Therefore, developing a method to identify promising proteins by predicting their performance based on their 3D structure is essential.

Recent advancements in machine learning, particularly AlphaFold2, have revolutionized the prediction of protein 3D structures^21–23^. While these models have demonstrated remarkable success in predicting the structures of natural proteins, their application to mutant proteins remains controversial^24,25^. This is partly because structural deviations caused by mutations can challenge the accuracy of current predictive models. Nonetheless, the intrinsic relationship between protein structure and function highlights the importance of incorporating 3D structural data into virtual screening approaches for protein design.

Many prior studies on virtual screening have primarily focused on treating protein sequences as linear strings, linking them to functional properties without explicitly incorporating 3D structural data. For example, Bedbrook et al.^16^ used Gaussian process regression to predict photo-induced current properties in channelrhodopsin (ChR) mutants based on their sequences. Similarly, Liu et al.^15^ employed deep learning to relate antibody sequences to their binding affinity for target proteins, aiding in the design of high-affinity antibodies.

In this study, we developed a machine learning framework, called ‘Foldinsight’ to predict the properties of target proteins from their amino acid sequences, utilizing AI-predicted structural information. This framework also offers the design of new amino acid sequences to improve protein functionality. To evaluate its utility, we applied the framework to a dataset of ChR variants^16^ along with their quantitative properties. Using AlphaFold2, we predicted the structures of the ChR variants and represented them as fixed-length energy state vectors, which were subsequently used for regression analysis to predict protein properties. This approach successfully predicted protein functionality and identified key structural regions essential for functionality. By doing so, it will be possible to better understand how mutations influence protein functionality, paving the way for more effective protein engineering strategies.

## Results

### Foldinsight: Framework to predict function from sequence via structure

Foldinsight is a machine learning framework, which predicts functionality of the target protein from amino acid sequences via AlphaFold2 and identifies key structural regions critical for function. The analysis proceeded as follows:

We began with a dataset consisting of amino acid sequences of the target protein variants and their corresponding experimentally measured functional values (Data Preparation, **Fig. 1A**). To incorporate 3D structural information, each amino acid sequence was used as input to AlphaFold2 for structure prediction (Structure Prediction; **Fig. 1B**). Since the predicted structures might not represent the most energetically stable conformations, we further refined them via energy minimization to produce more stable, relaxed structures (Structure Stabilization; **Fig. 1C**).

**Figure 1.**
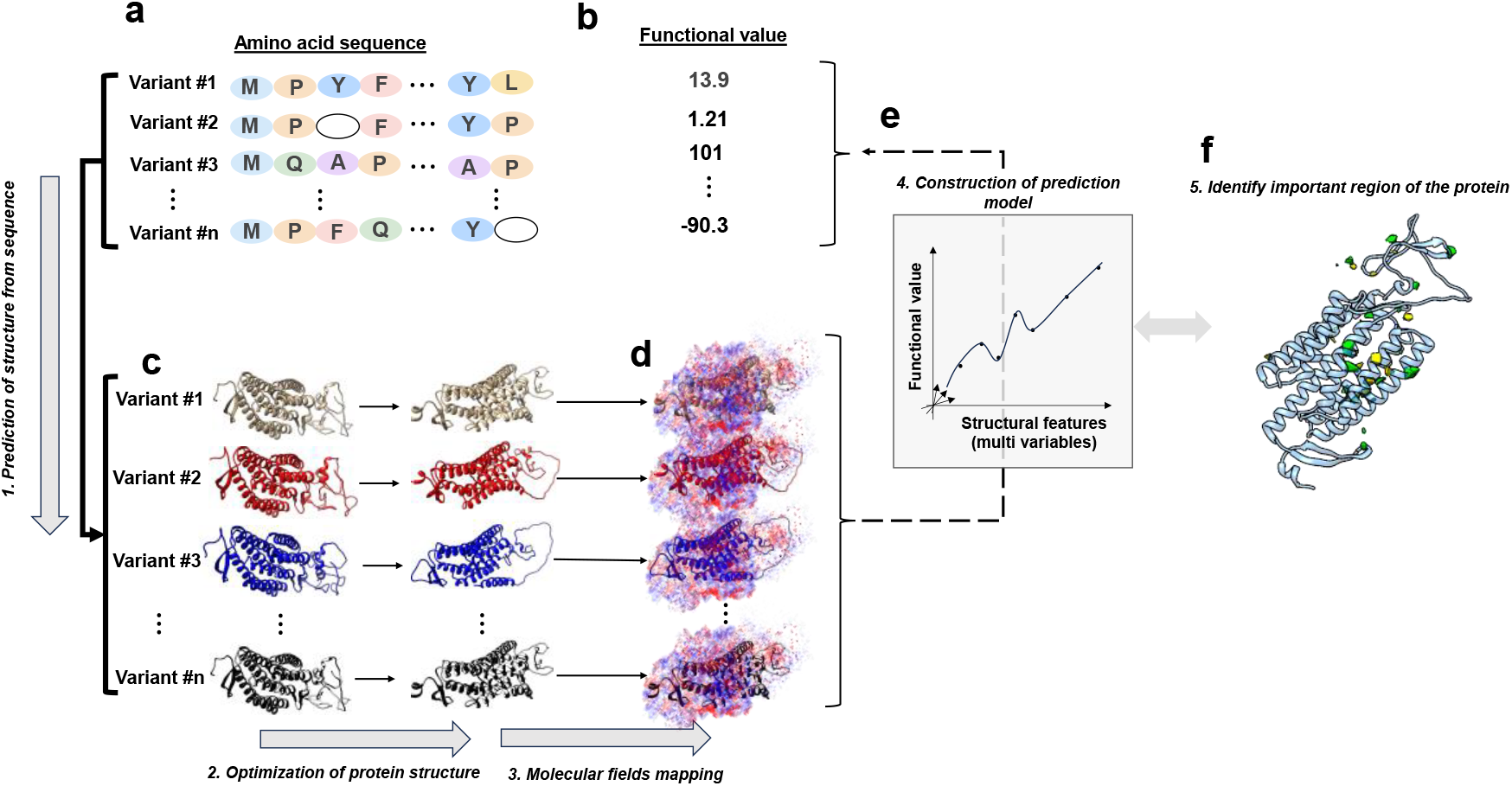
Machine learning framework for predicting protein functionality and identifying functionally important regions. **a** Amino acid sequences of protein variants were used as input to AlphaFold2 for structure prediction. **b** Each sequence is associated with functional value. **c** The predicted structures were further refined via energy minimization to obtain more stable conformations. **d** Molecular field mapping was applied to extract structural features, where a virtual probe atom was placed in a 3D lattice surrounding the protein to quantify van der Waals and Coulomb interactions. Red and blue colors represent the values of potential in lattice points. **e** These structural features were used as input variables for a regression model to predict protein functionality. **f** By analyzing the regression model, functionally important structural regions were identified.

Next, we extracted physicochemical features from the refined protein structures using a “molecular field” approach (Molecular Field Representation; **Fig. 1D**). Conceptually, this involved placing a virtual carbon probe throughout a three-dimensional lattice that enveloped the protein and quantifying the local intermolecular interactions—specifically, van der Waals and Coulomb forces—experienced by the probe. These interactions were then converted into a fixed-length vector for each protein variant.

The feature vectors derived from the molecular field were employed as input to predict the protein’s functional values by regression (Functional Prediction; **Fig. 1E**). By training relationships between protein variants and their measured functionalities, we were able to accurately predict the performance of new variants. Finally, by analyzing the regression coefficients and their influence on predicted functionality, we pinpointed particular structural regions that have a pronounced impact on protein performance (Identification of Functionally Important Regions, **Fig. 1F**). This approach not only predicts protein function, but also elucidates the underlying structure-function relationship.

### Dataset of ChR variants

We applied Foldinsight to a publicly available dataset of ChR variants reported by Bedbrook et al. ^16^ (**Table 1**). The dataset consists of 154 variants of amino acid sequence and their photo-induced current properties: max peak off-kinetics and green norm (**Fig. 2A**). Notably, the amino acid sequences differ in length (**Fig. 2B**), which can complicate the use of standard machine learning, which typically requires fixed-length inputs. Additionally, the mutation and deletion sites are relatively uniform in their distribution across the sequences (**Fig. 2C**). No clear correlations were observed among the three measured properties (**Fig. 2D**), suggesting that they may be governed by different local sequences or structures.

**Figure 2.**
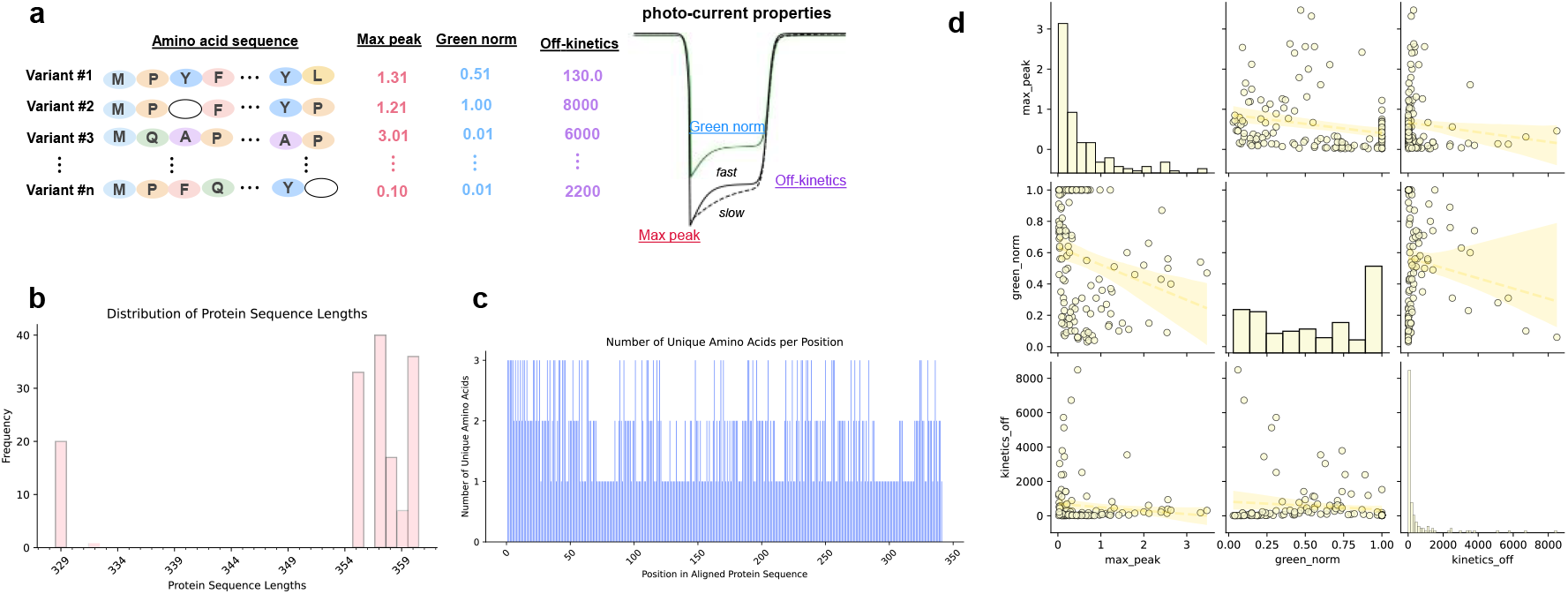
Overview of the dataset used in this study. **a** The dataset consists of 154 channelrhodopsin (ChR) variants with amino acid sequence differences and their experimentally measured photo-induced current properties: max peak, off-kinetics, and green norm. **b** Distribution of protein sequence lengths across the dataset. **c** Distribution of unique amino acid residues at each position in the aligned sequences. **d** Pairwise scatter plots of the three measured photo-current properties.

In prior work, multiple sequence alignment was performed to represent all variants as a fixed-length vector, where deleted missing amino acids are treated as mutations. They used Gaussian process regression to predict functionality directly from the aligned sequences without structural information. In order to compare our structure-based approach with this earlier sequence-based approach, we employed the exact same training data as used in the previous research.

### Protein structure by AlphaFold

To conduct a structure-based analysis, we used AlphaFold2 to convert each channelrhodopsin (ChR) sequence into a three-dimensional structure (**Fig. 3A**). To assess AlphaFold2’s performance, we confirmed that the predicted structures had plDDT scores above 70, indicating high reliability (**Fig. 3B**). Although AlphaFold2 excels at predicting wild-type protein structures, its performance on mutant proteins remains a point of debate (refs). Therefore, we further refined both side-chain and backbone positions by performing energy minimization in Chimera^26^. Next, to evaluate differences among the predicted 3D structures of each variant, we conducted a Canonical Correspondence Analysis (CCA) on the α-carbon coordinates of these refined ChR structures (**Fig. 3C**). These results indicate that α-carbons in close proximity exhibit strong correlation, whereas those that are far apart generally do not correlate but do show strong correlation at certain locations, which consists of our intuitive expectation.

**Figure 3.**
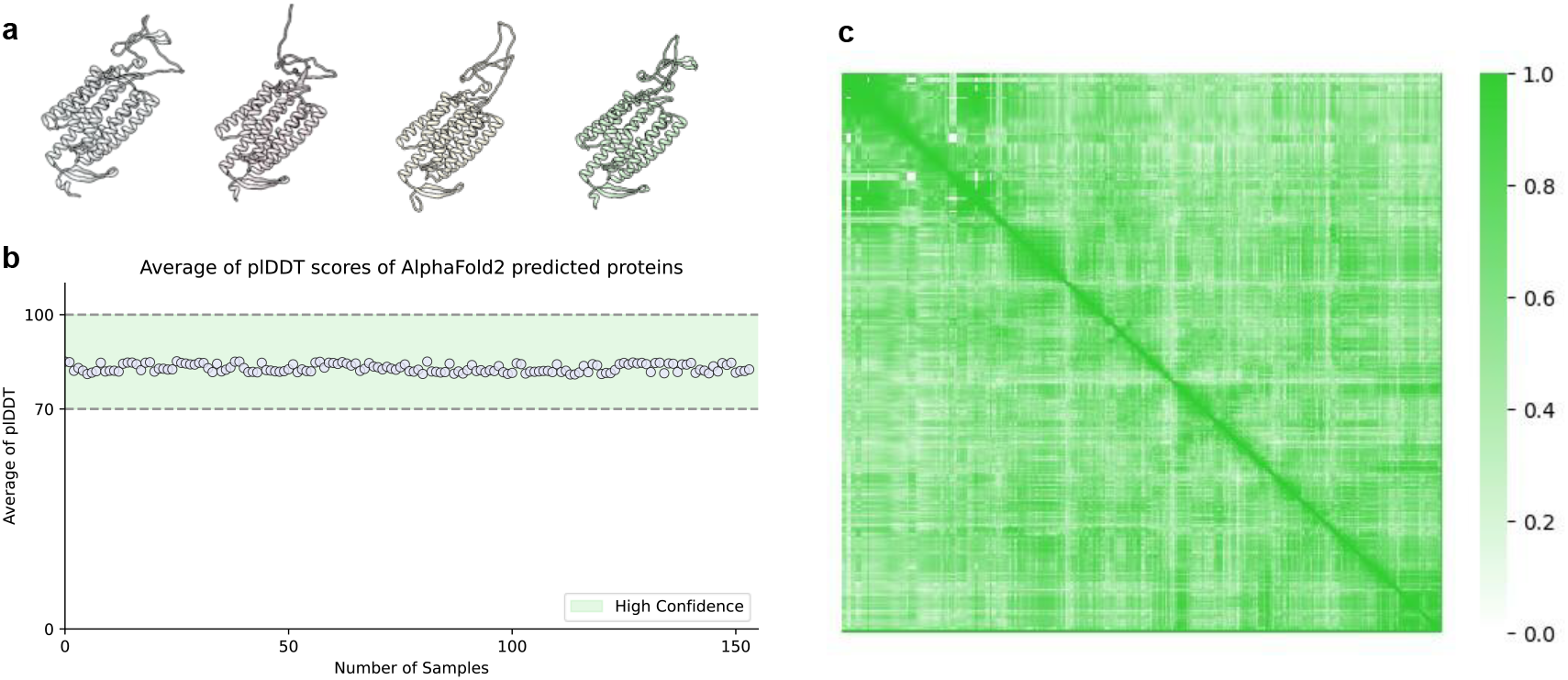
Structure-based analysis of channelrhodopsin (ChR) variants. **a** Representative 3D structures of ChR variants predicted by AlphaFold2. **b** pLDDT scores corresponding to all predicted structures. **c** Canonical Correspondence Analysis (CCA) of α-carbon coordinates in the refined ChR structures. The heatmap represents the correlation between α-carbon positions across the dataset.

### Fixed-length features of D structure

To represent the predicted 3D structures of channelrhodopsin (ChR) variants into fixed-length vectors, we employed a molecular field mapping approach (Ota et al.^29^, and CoMFA^37^). In this method, a probe atom (an sp^3^ carbon carrying a +1 charge) is placed at systematically arranged lattice points around the ChR structure. At each lattice point, both van der Waals and Coulomb potentials are calculated, generating molecular field values that are then transformed into fixed-length vectors. These vectors, representing the molecular fields, can be used as explanatory variables for predicting functionality of each ChR variant in our analyses. We then visualized these vectors by employing dimension reduction with PCA ^38^. In the PCA space, we observed a relationship between the reduced features and the performance values of the ChR variants (**Fig. 4**).

**Figure 4.**
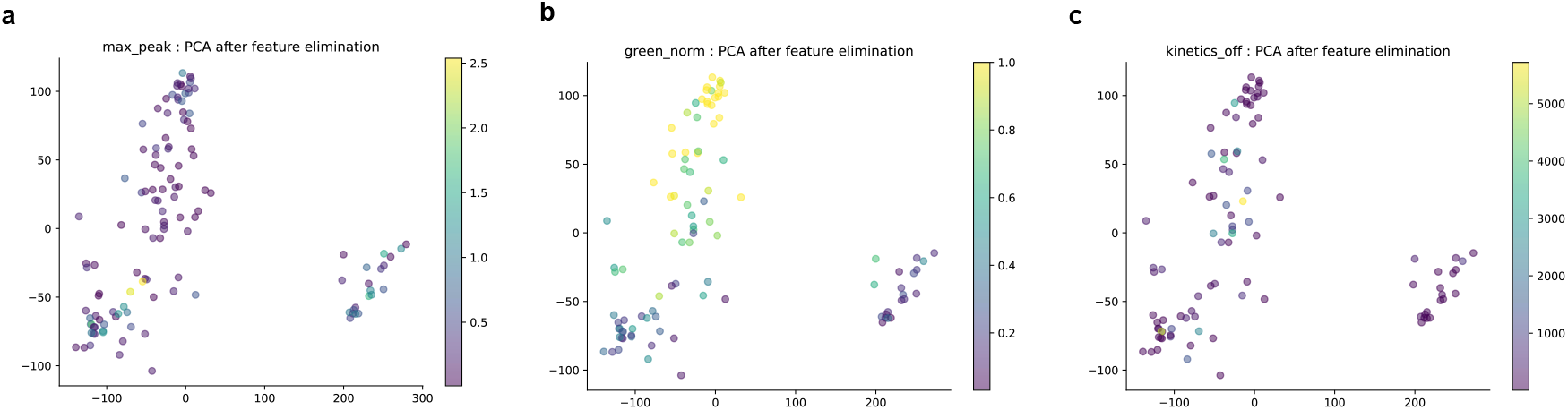
Principal component analysis PCA of molecular field representations. **a** PCA projection of molecular field features colored by max peak values. **b** PCA projection of molecular field features colored by green norm values. **c** PCA projection of molecular field features colored by off-kinetics values.

### ChR function prediction and its key regions

To predict the Max peak, Green norm, and Kinetics off values of ChR from the fixed-length vectors of the molecular field, we used Partial Least Squares Regression (PLS). As a result, the predicted values were correlated with experimental values (**Fig. 5A**), indicating that the AlphaFold2-predicted structure of target protein variants indeed contains information to predict their performances. In addition, we showed that the prediction accuracy of functional properties decreased in the absence of structural refinement by energy minimization after AlphaFold2 (**Fig. 5B**), suggesting that additional refinement is crucial to extract the structural characteristics for each variant of the target protein.

**Figure 5.**
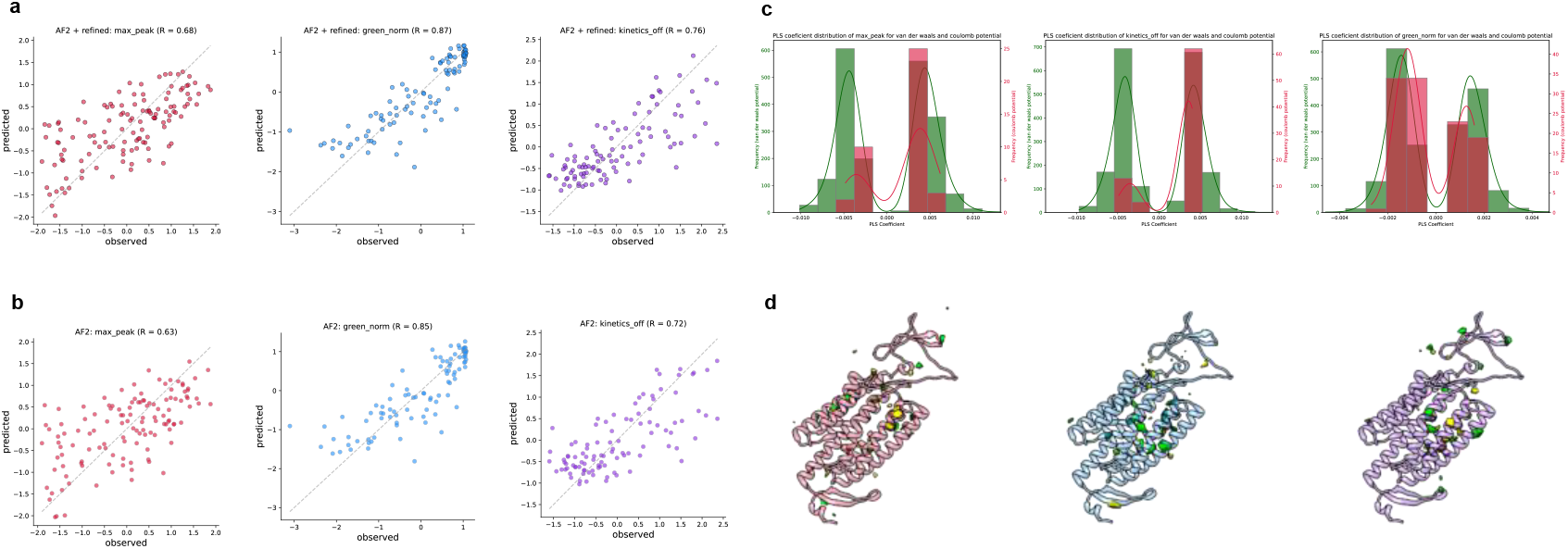
Prediction of ChR functional properties and identification of key structural regions. **a** Max peak, Green norm, and Kinetics off predicted using Partial Least Squares Regression (PLS) from the molecular field vectors. **b** Max peak, Green norm, and Kinetics off predicted using PLS in the absence of structural refinement via energy minimization after AlphaFold2. **c** Distributions of PLS regression coefficients for van der Waals and Coulomb potentials for Max peak, Green norm, and Kinetics off. **d** Contour map highlighting key regions in the ChR structure based on van der Waals potentials. The contour map is overlaid on the structure of C1C2, a representative ChR variant. Green regions indicate positive correlations with functional properties, whereas yellow regions indicate negative correlations.

Next, we examined which energies are important for predicting functional properties of ChR variants. By plotting distributions of regression coefficients of PLS, we found that van der Waals potentials have stronger weight than coulomb potentials (**Fig. 5C**). This result suggests that, in the function of the target protein, the spatial arrangement and density of atoms, which reflect van der Waals potentials, play a more critical role than partial charges or electrostatic interactions.

Finally, to identify key regions in the ChR structure using van der Waals potentials, we generated a contour map highlighting lattice points with larger (regardless of sign) PLS regression coefficients for van der Waals potentials (see Methods). **Fig. 5D** shows the key regions mapped onto a representative ChR structure. In this contour map, green regions indicate a positive correlation between van der Waals potentials and ChR’s functional properties, whereas yellow regions indicate a negative correlation. Because Van der Waals potentials are higher in spatially denser regions and lower in sparse regions, in green regions, substituting amino acids with larger side chains for those with smaller side chains may improve performance. Conversely, in yellow regions, substituting amino acids with smaller side chains for those with larger side chains is expected to enhance performance. In addition, we applied a similar analysis to Coulomb potentials; however, no distinct contour patterns were observed.

## Discussion

In this study, we developed a computational framework called Foldinsight for protein engineering that integrates three-dimensional protein structures. Protein sequences were converted into 3D structures using AlphaFold2 with further structural optimization and then transformed into fixed-length vectors through Molecular Field Mapping to preserve physicochemical properties. This reformulates the task of predicting protein properties into a standard regression problem using fixed-length descriptors. Applying the framework to public ChR data, we successfully identified key regions contributing to the ChR functionality.

Our framework, incorporating protein three-dimensional structures, naturally aligns with the principle that protein function depends on its structure. Beyond its conceptual validity, this approach offers practical advantages. Traditional sequence-based analyses often parameterize positions of the mutation in the sequence. However, positions that have never undergone experimental mutations remain unparameterized, making it difficult to predict the effects of introducing mutations at these sites. Our method overcomes this limitation by converting protein structures into their surrounding energy states. Consequently, even for residues without prior experimental mutations, hypothetical mutations can be simulated, and the changes in energy states can be captured to predict performance. Additionally, our framework is able to identify key regions contributing to protein properties, providing actionable insights for protein design, such as guiding where and what types of residues (e.g., large or small) should be introduced. These capabilities are particularly valuable for efficiently optimizing protein properties using limited experimental data.

Previous studies have already demonstrated the prediction of ChR functionality using sequence-based approaches. In this study, we utilized the exact same dataset as Bedbrook et al., enabling a direct comparison with their methods. Bedbrook et al. employed Gaussian process regression for sequence-based analysis (see Ref.), achieving predictive accuracies of 0.77, 0.79, and 0.90 for max peak, off-kinetics, and green norm, respectively. Our framework demonstrated comparable predictive performance (Figure 5A). However, unlike their study, our approach directly incorporates structural information, offering the conceptual soundness of three-dimensional structure integration. As a result, our framework not only achieves comparable accuracy but also enables the identification of key protein regions, providing greater utility for protein engineering applications.

Foldinsight uses AlphaFold2 to reconstruct the three-dimensional structures of protein variants, based on which it predicts their properties. However, the reliability of AlphaFold2 in predicting the structures of mutated proteins remains a topic of debate. For example, AlphaFold2 finds it challenging to accurately predict structure-disrupting mutations, such as those that impair proper folding^24^. On the other hand, AlphaFold2 has been shown to detect localized structural distortions caused by mutations involving one to three amino acid residues^25^. Since AlphaFold2 does not simulate the folding process itself^39,40^, it is possible to detect local structural differences in variants if the variants underwent a folding process similar to that of the wild type. In this context, predicting the structures of protein variants with more than ten residue differences, as examined in this study, should be considered a challenging task. Nevertheless, our framework has achieved high accuracy in predicting their properties by leveraging structure-based information. This suggests that the applicability of AlphaFold2 may extend beyond initial expectations.

While our framework demonstrates the potential of integrating three-dimensional structural information into machine learning for protein engineering, there still are certain challenges. One major limitation is the uncertainty in AlphaFold2’s reliability when predicting the structures of mutated proteins, particularly those with extensive mutations that destabilize the overall structure. In stable structures, the surrounding energy states of a specific pose can be calculated and used as representative parameters to capture structural characteristics. However, for unstable or highly dynamic structures^41,42^, a single pose may not sufficiently represent the overall structural features, leading to inaccuracies in molecular field mapping and subsequent functional predictions. Moreover, the structural refinement process primarily relies on energy minimization in a simplified vacuum state^26^, which does not account for physiological conditions such as membrane-bound states^43,44^ or interactions with surrounding molecules^45^. Thus, for future study, enhancing prediction and refinement methods taking account for physiological factors will be crucial for overcoming these challenges and expanding the framework’s applicability.

## Methods

### Data collection

Channelrhodopsin (ChR) variant data were obtained from the study by Bedbrook et al.^16^, which includes the primary protein sequences and photo-induced current properties of various ChR variants. The properties recorded were peak photocurrent strength, wavelength sensitivity, and photocurrent decay rate, referred to as current strength, green norm, and off-kinetics, respectively.

### Data preprocessing

We predicted the structures of each ChR variant using Alphafold2 (version 2.1.0)^21,22^. Alphafold2 generated several alternative structures for each variant, and we selected the best structure based on the predicted Local Distance Difference Test (plDDT) score.

### Refinement of predicted ChR structures

To further optimize the predicted structures of the ChR variants, energy minimization was performed using the “Minimize Structure” tool in UCSF Chimera (version 1.16)^26^. The “Steepest descent steps” and “Conjugate gradient steps” were set to 1000 and 100, respectively, while other parameters were left at their default settings.

### Structural Alignment and Superimposition of ChR ariants

To ensure consistency in the orientation of all predicted ChR variant structures, structural alignment was performed by modeller (version 10.4)^27^, and they were superimposed in the same orientation.

### Canonical Correlation Analysis

To investigate the internal structural correlations among mutant proteins, CCA (Canonical Correlation Analysis)^28^ was employed. After the structures of the variants were predicted using AlphaFold2, pairwise positional vector data of all combinations of α-carbon atoms were prepared and analyzed using CCA. The correlation coefficient of the first canonical component output by CCA was evaluated as the measure of internal structural correlation.

### Molecular field mapping

To convert the three-dimensional structures of ChR variants into fixed-length numerical vectors, molecular field mapping was conducted^29^. Molecular field mapping quantifies energies such as the van der Waals and Coulomb potentials around a molecule. In this method, the ChR was enclosed in a 1 Å lattice, and probe atoms were placed at each lattice point. The probe atom was assumed to have the van der Waals property of a sp3 carbon and a charge of +1. Then, the interaction energy (van der Waals potential and Coulomb potential) between the probe atoms and each atom of the ChR was summed and assigned to each lattice point. The van der Waals potential and Coulomb potential were calculated according to the following equations^30,31^.

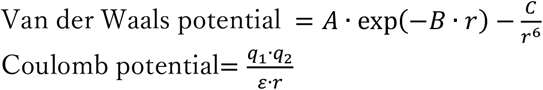

The parameters A, B, and C in the van der Waals potential’s equation are fixed values that depend on the combination of interacting atoms, and their values were taken from the paper by Vinter et al^31^. The parameters q1 and q2 in the Coulomb potential’s equation represent the charges of each atom, and epsilon is the dielectric constant. The variable r represents the distance between the atoms in both equations.

### Construction of prediction models

The overall learning and evaluation framework in this study was similar to that used by Bedbrook et al, who originally collected the data. Following this framework, we performed a 20-fold cross-validation, aggregated all test data across the folds and evaluated the models using the Pearson correlation coefficient between the measured and predicted values.

### Feature selection

To avoid data leakage^32^, feature selection was performed only on the training folds. In the calculation of energies, the values could diverge when the atoms on the protein and the probe atoms were extremely close. To mitigate the effect of these outliers, the upper and lower bounds of the energy values were set to the 95th and 5th percentiles of the dataset, respectively. Moreover, for each lattice, features with high skewness (greater than 2.5) were excluded to improve the model stability. Additionally, features with low variance (less than 2 kcal) were also excluded. To further refine the feature set, recursive feature elimination (RFE)^33^ was performed following the initial filtering. Feature importance during the RFE process was determined using a linear support vector regression model. The final number of selected features was set to 1500. As a result, the dimension was reduced from about 1,000,000 to 1,500.

### Machine learning model

To construct the predictive model, partial least squares (PLS) regression^34^ was performed. The optimal number of dimensions, which is a hyperparameter of PLS, was selected based on the highest average Pearson correlation coefficient obtained through 5-fold cross-validation. This cross-validation was performed within the training folds of the 20-fold cross-validation of the entire dataset (i.e., double cross-validation approach^35^).

### Visualization of structural importance

Visualization of key regions with a significant influence on photo-induced current properties was performed as follows: using PLS (partial least squares) modeling, the importance of each grid point was calculated as a weight (PLS partial regression coefficient times variance on each lattice point), and based on this weight, the regions most sensitive to the properties were identified. The grid points selected as features were connected by the Delaunay algorithm^36^, and weights were linearly interpolated on the edges between each grid point. This resulted in the computation of endpoints corresponding to the “top 99% and bottom 1% weights.” The calculated endpoints were connected with distances of 5 Å or less as nearest neighbors, and the area encompassed by the resulting connected endpoints was defined as the key region. Chimera (version 1.16) was used for the visualization.

## Acknowledgment

This work was partly supported by the Japan Agency for Medical Research and Development (AMED; grant numbers JP21wm0525004; to M.S., H.N., R.K. and JP24wm0625416; to Y.Y., O.K.), JSPS KAKENHI (grant number JP24K20893; to R.O.), and the Moonshot R&D– MILLENNIA Program (grant number JPMJMS2024-9; to H.N.). Computations were partially performed on the NIG supercomputer at ROIS National Institute of Genetics. The authors thank Dr. Ryota Kobayashi at the University of Tokyo for his invaluable advice during our discussions. The authors are grateful to the members of the Data-driven biology Laboratory for their helpful input.

## Author Contributions

R.O., M.S., W.A. and H.N. conceived the project. R.O. developed the model and analyzed the data. R.O. and H.N. wrote the manuscript with input from all the authors.

## DATA AVAILABILITY

Datasets for the current study are obtained from the paper (doi:10.1038/s41592-019-0583-8)

## CODE AVAILABILITY

All code used in the current study will be available on the GitHub repository after publication.

